# Using Polycistronic gRNAs and a tRNA Cleavage System in CRISPR-Based Genome Editing of *Toxoplasma gondii*

**DOI:** 10.64898/2026.10.12.755625

**Authors:** Janessa Grech, Melanie Teng, Renfu Wang, Matthew Child

## Abstract

CRISPR/Cas9 is widely used for genetic manipulation in *Toxoplasma gondii*, but optimizing gRNA expression and multiplexed editing remains challenging. Here, we evaluated alternative promoters, gRNA scaffolds, and a tRNA-based polycistronic system to improve editing efficiency and reduce synthetic DNA costs. Four constructs targeting the SAG1 locus were used to compare the U6 and H1 promoters, as well as standard and modified gRNA scaffolds. U6-driven constructs supported robust gRNA-targeted Cas9 activity, whereas H1-driven constructs failed to produce detectable phenotypes. The alternative gRNA scaffold, modified by a number of base changes and insertions, increased knockout efficiency compared to the standard scaffold. To simplify dual gRNA-based methods, we establish a tRNA-based polycistronic system to simultaneously target two independent loci. Targeting the SAG1 and UPRT locus for proof-of-concept, all U6-driven polycistronic constructs conferred efficient SAG1 knockout and 5-fluorodeoxyuridine resistance, whereas H1-driven constructs exhibited position-dependent activity, with only the 3′ gRNA being functional. These results demonstrate that U6-driven, tRNA-based polycistronic gRNA expression enables reliable multiplexed genome editing in *T. gondii*, with the minimal H1 promoter being insufficient for consistent multi-gRNA activity. This optimized system provides a scalable framework for dual-gRNA CRISPR strategies in this organism.

## Introduction

CRISPR/Cas systems have revolutionized genome editing by providing a versatile and efficient means of introducing targeted genetic modifications. A key advancement in increasing the complexity of CRISPR applications is the development of polycistronic guide RNA (gRNA) expression strategies to increase multiplexing potential^1–4^.

Polycistronic gRNA expression strategies involve the contiguous expression of multiple gRNAs from a single transcript. One effective approach for processing these polycistronic gRNAs involves incorporating transfer RNA (tRNA) sequences as spacers between individual gRNA units^1,2^. This approach leverages the endogenous tRNA processing machinery (RNase P and Z) to cleave precursor tRNAs at defined sites^5^. By flanking each gRNA with tRNA sequences, the transcript is accurately processed into individual, functional gRNAs *in vivo*^2^. This method has been reported to maintain high expression levels relative to individual gRNAs and has been successfully applied in plants and mammalian cells^1,2,4^.

While CRISPR/Cas9 systems are used for targeted genome editing in *T. gondii*, and is well-established for gene disruption, modification, and screening^6–13^, there is no published evidence demonstrating the use of polycistronic gRNA systems for the contiguous expression of multiple gRNAs from a single transcript. Given *T. gondii*’s amenability to CRISPR-based manipulation, and the increasing demand for high-throughput high-complexity functional genomics, adapting the tRNA-gRNA strategy could offer a promising approach for multi-site editing in this model apicomplexan. Here, we sought to expand the CRISPR/Cas9-mediated genome editing toolbox for *T. gondii* by testing both single and arrayed gRNA expression, targeting the SAG1 and UPRT loci. To further enhance editing efficiency we sought to reduce construct size and remove repetitive elements that can be a challenge during plasmid construction. Finally, we evaluated promoters^14^ and gRNA scaffolds^15^ alternative to the ones typically used in this organism^6,11^.

## Results

### Optimization of Cas9 in *Toxoplasma gondii*

The U6 promoter has been the go-to promoter to drive gRNA expression in *T. gondii*, providing robust editing efficiency^6,9–11^. However, its relatively large size increases the cost of synthetic DNA constructs, and alternative promoters may offer advantages in transfection efficiency and vector design. Similarly, gRNA scaffold modifications have the potential to enhance editing efficiency^15^. Therefore, despite the widespread use of CRISPR/Cas9 technology, challenges remain in optimizing gRNA expression, improving transfection efficiency, and enabling multi-site editing in *T. gondii*. For this reason, our primary aim was to optimise the CRISPR/Cas9 system in this apicomplexan parasite for facile cloning, improved transfection efficiency, and reduced cost of synthetic DNA fragment production. To this end, we first evaluated alternative promoters and gRNA scaffolds.

Four gRNA expression cassettes were generated, all targeting the major surface antigen SAG1. SAG1 was chosen to evaluate on-target Cas9 activity since its knockout gives a clear readout via immunofluorescence, and a functional gRNA sequence has already been previously described by Sidik *et al.*^6 10^To assess whether a shorter promoter could replace the standard U6 promoter, we tested the H1 promoter^14^. The U6 promoter, first adapted for use in *T. gondii* by Sidik et al.^6^, is around 440 base pairs in length, whereas H1 promoter is approximately 100 bp. As for evaluating the efficiency of a possible alternative gRNA scaffold, the standard scaffold, here referred to as scaffold 1, was compared to a modified version (cr2 or scaffold 2)^15^.

Knockout efficiency for the different constructs was assessed by quantifying parasitophorous vacuoles negative for SAG1 staining (Fig. 2a). Significant differences were observed between constructs utilising the H1 and U6 promoters, as well as between those expressing the standard and modified gRNA scaffolds (Fig. 2b). Both U6-driven constructs resulted in efficient SAG1 disruption, whereas neither H1-driven construct produced a detectable phenotypic change. Notably, the construct incorporating the modified gRNA scaffold yielded a significantly higher proportion of SAG1 knockout compared with the standard scaffold, the latter of which resulted in a knockout efficiency comparable to that previously published^6^.

**Figure 1.**
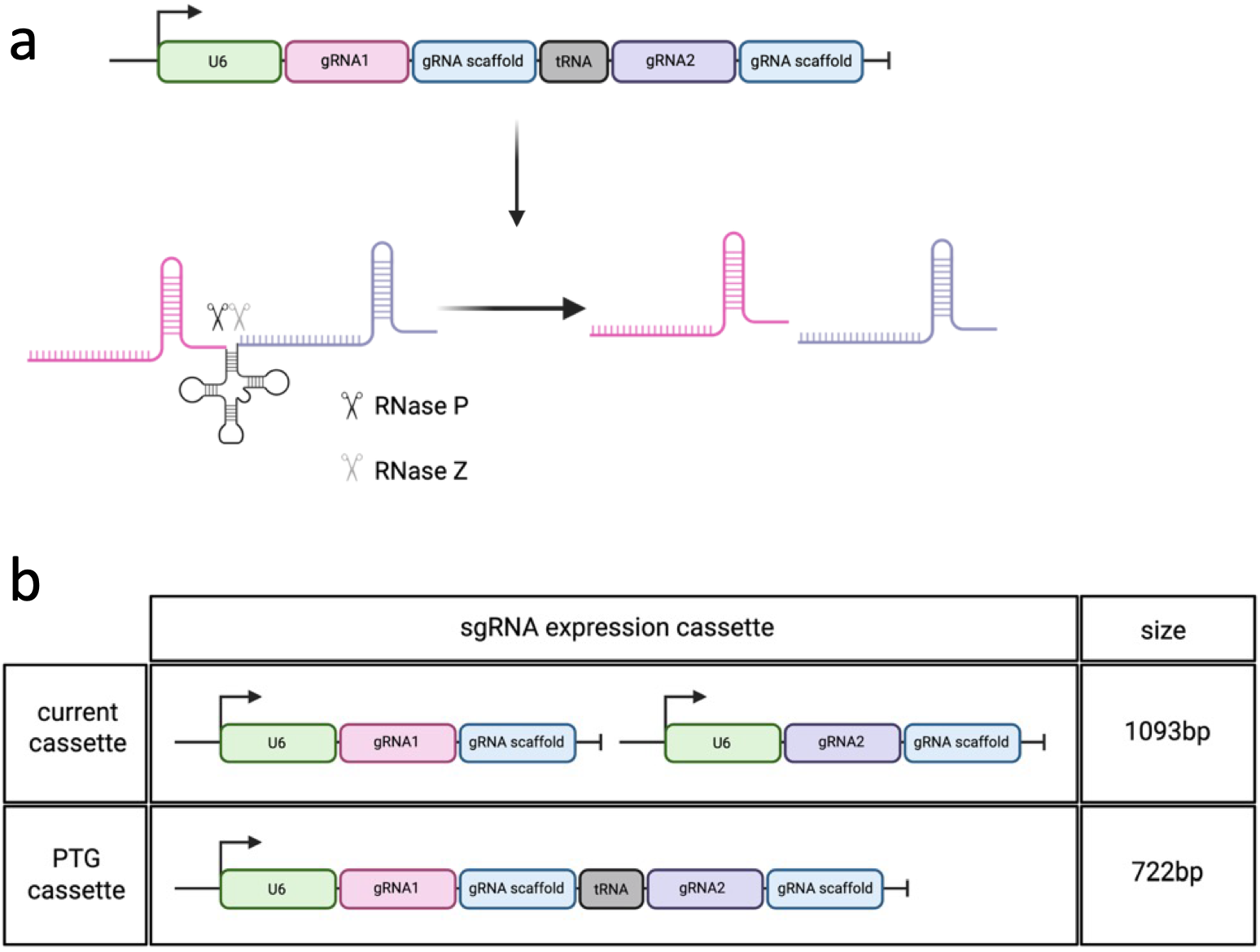
Polycistronic tRNA–gRNA (PTG) strategy for multiplex CRISPR guide RNA expression. **(a)** Schematic illustration of the PTG system for expressing two single guide RNAs using a single RNA polymerase III promoter. The endogenous tRNA-processing machinery (RNase P and RNase Z) recognizes the tRNA structure and precisely cleaves the transcript to release individual mature gRNAs containing intact scaffold sequences. **(b)** In the conventional configuration (current cassette), each gRNA is independently transcribed under separate U6 promoters, thus requiring a minimal total cassette size of 1093 bp. In the PTG design, a single U6 promoter drives a polycistronic transcript in which gRNA1 and gRNA2 are separated by a tRNA sequence. This configuration reduces cassette size to 722 bp while enabling efficient multiplex genome editing.

**Figure 2.**
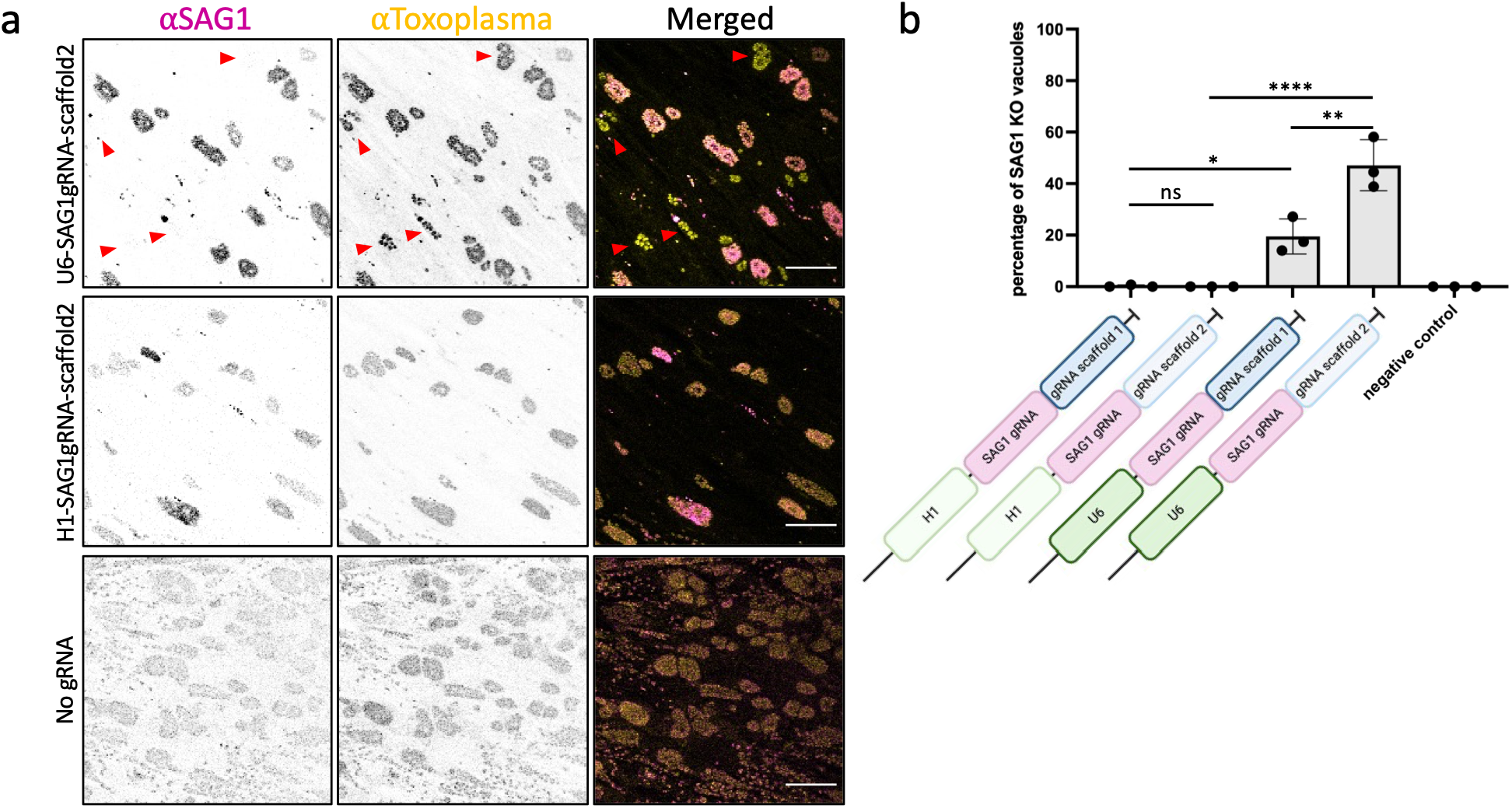
Single-gRNA editing efficiency at the SAG1 locus. **(a)** Representative immunofluorescence images of parasitophorous vacuoles stained using anti-Toxoplasma antibody (yellow) and anti-SAG1 (magenta). Red arrows indicate parasitophorous vacuoles lacking detectable SAG1 staining. Scale bar: 50μm. **(b)** Quantification of SAG1 knockout efficiency for each construct. Knockout efficiency was calculated as the percentage of vacuoles lacking detectable SAG1 staining. Data represent the mean ± SD of three biological replicates, with ≥100 vacuoles counted per replicate. Statistical comparisons were performed using one-way ANOVA with Tukey’s post hoc test. ns (P > 0.05), * (P ≤ 0.05), ** (P ≤ 0.01), *** (P ≤ 0.001), **** (P ≤ 0.0001)

### Implementation of a tRNA-based polycistronic gRNA system

Next, we investigated whether a tRNA-based polycistronic system could be used to express arrayed gRNAs in *T. gondii,* similar to other systems. Six constructs were designed to target two independent loci using a single contiguous gRNA array cassette. We compared the use of human and *T. gondii* glycine tRNAs while also assessing gRNA activity at different positions within the polycistronic transcript. The promoter and gRNA scaffold variants were also incorporated into the polycistronic expression cassettes. The two loci targeted by the six constructs designed were the SAG1 locus and the UPRT locus. The UPRT gene encodes uracil phosphoribosyltransferase, a key enzyme in the pyrimidine salvage pathway, the disruption of which confers resistance to 5-fluorodeoxyuridine (FUDR)^16^.

As expected, none of the constructs expressing a single gRNA targeting SAG1 alone conferred resistance to FUDR (Fig. 3a). In contrast, all constructs expressing polycistronic gRNAs under the U6 promoter conferred FUDR resistance, regardless of the tRNA species used or the position of the gRNA within the polycistronic sequence. While constructs driven by the H1 promoter showed evidence of transcript expression, only the gRNA located at the 3′ end of the polycistronic sequence exhibited detectable activity. Although the reason for this was not fully investigated, it is possible that this expression was driven by the tRNA gene itself^17,18^.

**Figure 3.**
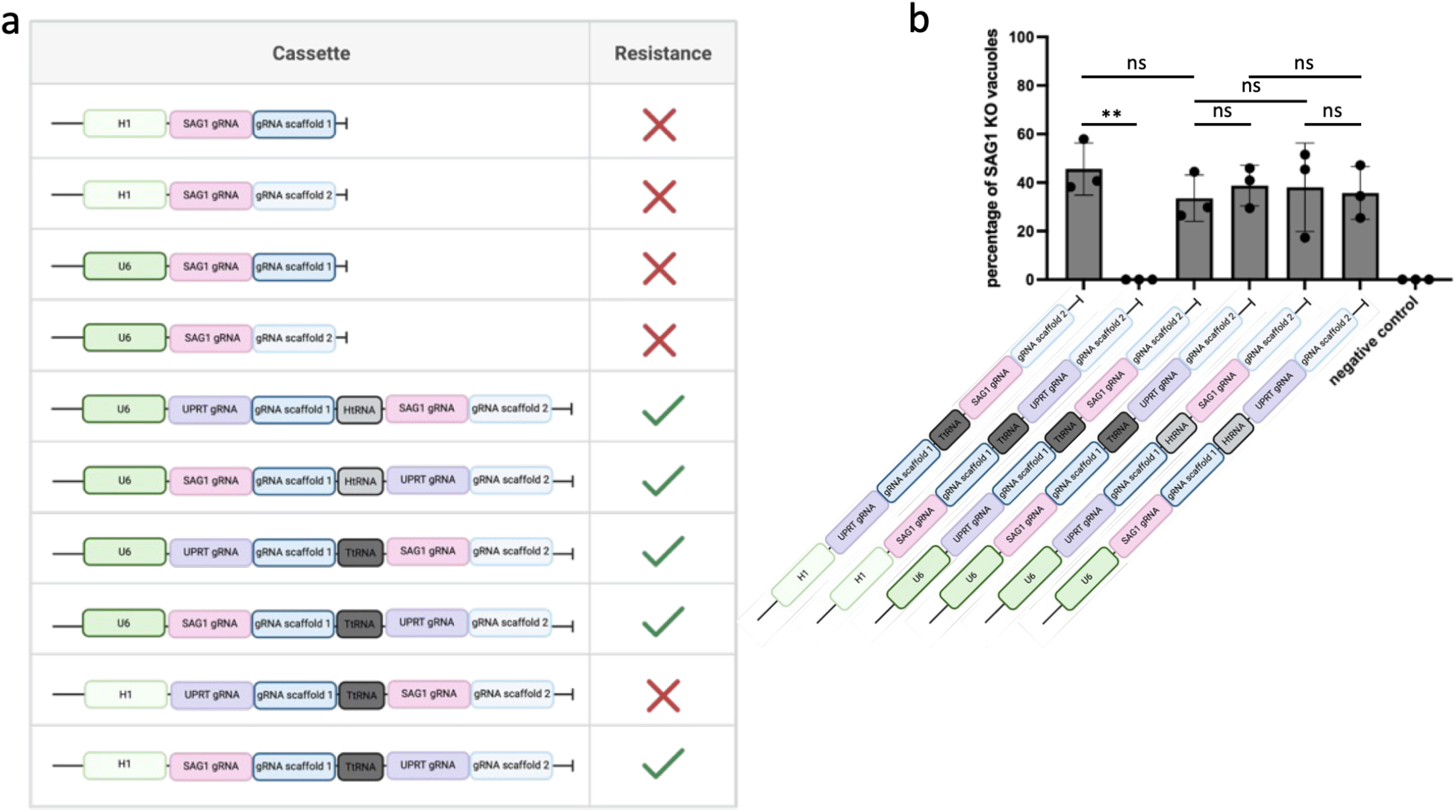
Polycistronic tRNA-based gRNA expression for multiplexed genome editing. **(a)** Resistance to 5-fluorodeoxyuridine (FUDR) in parasites expressing single or polycistronic gRNAs targeting SAG1 and UPRT loci. Constructs expressing the gRNA targeting the UPRT locus driven by the U6 promoter conferred FUDR resistance regardless of tRNA origin or gRNA position, whereas H1-driven constructs exhibited activity only for gRNAs at the 3′-end of the construct. **(b)** Quantification of SAG1 knockout efficiency for polycistronic constructs. Data represent the mean ± SD of three biological replicates, with ≥100 vacuoles counted per replicate. Statistical significance was assessed using one-way ANOVA with Tukey’s post hoc test. ns (P > 0.05), * (P ≤ 0.05), ** (P ≤ 0.01), *** (P ≤ 0.001), **** (P ≤ 0.0001)

We next sought to quantify knockout efficiency by determining the percentage of parasitophorous vacuoles negative for SAG1. All four U6-driven polycistronic constructs resulted in SAG1 knockout at efficiencies comparable to those observed with the single-gRNA system (P > 0.2) (Fig. 2b, Fig. 3b), indicating that multiplexed gRNA expression does not compromise editing efficiency when expressed under the U6 promoter. H1-driven polycistronic constructs again exhibited position-dependent activity, with only the 3′-end gRNA being functional. This is consistent with the observation that parasites expressing the UPRT-targeting gRNA positioned at the 5′ end of the polycistronic transcript failed to survive drug selection. All together, these results suggest that the reduced activity in H1 constructs is promoter-dependent rather than due to inefficient tRNA-mediated processing.

## Discussion

CRISPR/Cas9 technology has become a powerful tool for genetic manipulation in *Toxoplasma gondii*, enabling gene disruption, epitope tagging, and genome-wide functional screening^6,8–13^. Despite these advances, approaches for efficient multi-site genome editing remain largely unexplored. In other systems, polycistronic gRNA expression strategies have expanded the complexity of CRISPR-based editing strategies by enabling the contiguous expression of multiple independent gRNAs from a single polycistronic transcript. The tRNA-gRNA strategy in particular, has been successfully applied in plant and mammalian cells and supports efficient multi-site editing while maintaining high expression levels^1,2,4^.

In this study, we sought to expand the repertoire of tools available for use in *T. gondii*. By testing out a new promoter^14^, gRNA scaffold^15^, and an array-based gRNA system^2^, we aimed to reduce the size and complexity of dual-guide systems, thus reducing cost and increasing efficiency and usability.

The initial experiment aimed to reduce the size of the promoter from 444bp (U6) to 101bp (H1). We demonstrated that single gRNA expression driven by the U6 promoter consistently supported efficient genome editing in *T. gondii*, whereas the shorter H1 failed to result in any CRISPR-induced knockouts. In contrast, our attempt to investigate the use of an alternate gRNA scaffold showed that the construct with the modified scaffold not only resulted in successful genome editing but was significantly more efficient than that traditionally used, effectively doubling the frequency of knockouts generated using the SAG1 gRNA. This finding is noteworthy, particularly within the context of CRISPR-based experiments, as a significant increase in Cas9-mediated cleavage will undoubtedly improve cost-effectiveness and ultimately, experimental success.

To further improve genome editing in *T. gondii*, we endeavoured to adapt a tRNA-based polycistronic system for use in this parasite. While doing so, we again tested both promoters and tracrRNA scaffolds, as well as *T. gondii* and human tRNA sequences. Our results showed that the arrayed gRNA system enabled effective multi-site genome editing when driven by the U6 promoter, with no detectable reduction in editing efficiency at the SAG1 locus compared with single-gRNA constructs. These findings suggest that tRNA-mediated processing of polycistronic transcripts is functional in *T. gondii,* and the efficiency of genome editing is largely independent of the tRNA species of origin or gRNA position within the construct when promoter activity is adequate. It was also observed that H1-driven constructs exhibited position-dependent gRNA activity, with the gRNAs immediately downstream of the promoter seemingly being non-functional. This suggested that the issue does not lie with the tRNA-based polycistronic system but with the H1 promoter itself.

Together, these results highlight the critical role of promoter selection for both single and multi-site CRISPR/Cas9 applications in *T. gondii*. A U6-driven, tRNA-based polycistronic gRNA system provides a robust and scalable approach for multi-site genome editing, whereas the H1 promoter is not suitable for expression of arrayed-gRNAs. While the mechanisms underlying reduced activity in H1-driven polycistronic constructs were not directly assessed, future studies incorporating broader target validation and transcript analysis will help clarify these effects. It is also worth noting that the H1 promoter was adapted from the RNA component of human RNase P^14,19^, and though RNase P has been investigated in *T. gondii*, the RNA component has not been identified yet^21^. It is entirely possible that were the *T. gondii*-equivalent promoter were to be used, fully functional transcripts could be produced.

## Conclusion

In summary, the use of polycistronic gRNAs coupled with tRNA-based cleavage systems represents a powerful tool for complex genome engineering tasks. The tRNA-gRNA architecture offers several advantages; it is compact, promoter-efficient, and compatible with various CRISPR systems, including Cas9 and Cas12a^20^. Importantly, this system ensures precise and reliable processing without the need for additional exogenous enzymes. By using conserved tRNA motifs, this method is well-suited for use in both prokaryotic and eukaryotic organisms, and has demonstrated broad compatibility across species^1,2,22,23^. As CRISPR applications continue to expand, this strategy provides an elegant solution for multiplexed editing, synthetic biology, and large-scale functional genomics studies.

## Materials and Methods

### Plasmid design and construction

Four single-gRNA expression constructs targeting the SAG1 locus were generated to compare the U6 (440 bp) and H1 (∼110 bp) promoters^6,11,15^, as well as the standard and modified gRNA scaffolds^6,11,14^. The gRNA sequences were based on those described by Benns et al.^10^ and Sidik et al.^6^. Six constructs were also designed to trial the use of the tRNA-based polycistronic gRNA strategy, these constructs simultaneously targeting the SAG1 and UPRT loci. Both human and *T. gondii* glycine tRNAs were used to separate gRNAs within the transcript. gRNAs were positioned at different locations within the polycistronic sequence to assess position-dependent activity. Both U6 and H1 promoters, as well as standard and modified gRNA scaffolds, were incorporated into these constructs.

All constructs (Suppl. Table 1) were synthesised by Twist Bioscience as fragments. These were amplified by PCR using Q5® Hot Start High-Fidelity DNA Polymerase (NEB). The pCORe plasmid containing a Cas9 expression cassette and an ampicillin selection marker was used as the vector plasmid^10^. This was initially digested by BsaI-HF®v2 (NEB) to remove the mScarlett construct, after which the fragments were inserted via Gibson assembly (NEB Gibson Assembly® Cloning Kit). The plasmids constructed were transformed by heat shock and amplified using DH5α chemically competent *E. coli* (NEB) and extracted using the Monarch® Spin Plasmid Miniprep Kit (NEB). Sequencing to confirm correct plasmid construction was performed by Azenta.

### Cell culture

Human foreskin fibroblasts (HFF) obtained from ATCC (SCRC-1041™) were grown in Dulbecco’s modified Eagle’s medium (DMEM, Sigma D6429) supplemented with 10% fetal calf serum (FCS, Sigma F7524) and 2 mM L-glutamine (Gibco 25030). Cell culture was maintained at 37°C and 5% CO^2^. *Toxoplasma gondii* RH tachyzoites were grown on a monolayer of HFF cells under the same conditions until complete lysis of the monolayer, after which the parasites were transferred onto a new fresh monolayer for maintenance. Any parasite culture which was observed to be crashing, such as those undergoing drug selection, were egressed manually by scratching and syringing using a 26G needle and passaged onto new fresh HFF cells.

### Transfections

Parasites (10^5^) were transfected with 7 µg of plasmid DNA in 16-well Nucleocuvette strips using the Amaxa 4D-Nucleofector system (Lonza). The transfections were done in P3 Primary Cell 4D-Nucleofector® X Kit buffer (Lonza) using the program ‘FI-115’, after which the parasites were mixed with supplemented DMEM for recovery. Half of the resulting volume was used to infect coverslips seeded with HFFs, while the rest was used to infect T25 flasks for drug selection. Drug selection was initiated at 24-hours post-transfection using 5mM 5-fluorodeoxyuridine (Merck) and maintained for 7 days.

### Immunofluorescence Assays

Cells were fixed with ice-cold methanol, after which the parasites were stained to allow for the quantification of SAG1 knockouts. The cells were blocked and permeabilised using 3% BSA (Sigma) and 0.2% Triton X-100 (Sigma) in PBS (Sigma, D8537) for 1 hour at room temperature. Primary antibody incubation was done for an hour using anti-SAG1 (Invitrogen, MA5-18268) and anti-IMC12 (a generous gift from Prof. P. Bradley) or anti-Toxoplasma (Invitrogen, PA1-7252) antibodies. Secondary antibody incubation was done for one hour in the dark using anti-rabbit Alexa488 (Invitrogen, A11008) and anti-mouse Alexa594 (Invitrogen, A11032). The coverslips were mounted using Fluoromount-G™ (Invitrogen).

### Microscopy and image processing

Confocal microscopy was done using the Leica STELLARIS 5 inverted confocal microscope equipped with a HC PL APO CS2 20x/0.75 dry objective lens and DFC9000 sCMOS camera. Images were processed using Fiji. Confocal microscopy images were processed with Leica Application Suite X (LAS X) and Fiji (ImageJ version 2.14.0)^24^.

### Quantification and statistical analysis

SAG1 knockout efficiency was calculated as the percentage of parasitophorous vacuoles lacking detectable SAG1 staining. All experiments were performed in biological triplicate, with at least 100 vacuoles counted per replicate. Statistical analysis was done using GraphPad Prism, with significance between groups being assessed using one-way ANOVA with post hoc Tukey’s test for multiple comparisons. P-values less than 0.05 were considered significant. All statistically significant results are annotated with a line and asterisk(s) in the graphs.

## Acknowledgements

This work was supported by grants BB/M011178/1 and 202553/Z/16/Z from the Wellcome Trust (to M.A.C. and J.G respectively). The Facility for Imaging by Light Microscopy (FILM) at Imperial College London is part-supported by funding from the BBSRC (grant BB/T017929/1). Figure schematics were created using BioRender.com.

## References

1. Dong, F., Xie, K., Chen, Y., Yang, Y., and Mao, Y. (2017). Polycistronic tRNA and CRISPR guide-RNA enables highly efficient multiplexed genome engineering in human cells. Biochem. Biophys. Res. Commun. 482, 889–895. 10.1016/j.bbrc.2016.11.129.

2. Xie, K., Minkenberg, B., and Yang, Y. (2015). Boosting CRISPR/Cas9 multiplex editing capability with the endogenous tRNA-processing system. Proceedings of the National Academy of Sciences 112, 3570–3575. 10.1073/pnas.1420294112.

3. Zetsche, B., Heidenreich, M., Mohanraju, P., Fedorova, I., Kneppers, J., DeGennaro, E.M., Winblad, N., Choudhury, S.R., Abudayyeh, O.O., Gootenberg, J.S., et al. (2017). Multiplex gene editing by CRISPR–Cpf1 using a single crRNA array. Nat. Biotechnol. 35, 31–34. 10.1038/nbt.3737.

4. Knapp, D.J.H.F., Michaels, Y.S., Jamilly, M., Ferry, Q.R. V, Barbosa, H., Milne, T.A., and Fulga, T.A. (2019). Decoupling tRNA promoter and processing activities enables specific Pol-II Cas9 guide RNA expression. Nat. Commun. 10, 1490. 10.1038/s41467-019-09148-3.

5. Elder, J.J.H., Papadopoulos, R., Hayne, C.K., and Stanley, R.E. (2024). The making and breaking of tRNAs by ribonucleases. Trends in Genetics 40, 511–525. 10.1016/j.tig.2024.03.007.

6. Sidik, S.M., Hackett, C.G., Tran, F., Westwood, N.J., and Lourido, S. (2014). Efficient Genome Engineering of Toxoplasma gondii Using CRISPR/Cas9. PLoS One 9, e100450.

7. Markus, B.M., Bell, G.W., Lorenzi, H.A., and Lourido, S. (2019). Optimizing Systems for Cas9 Expression in Toxoplasma gondii. mSphere 4, 10.1128/msphere.00386-19. 10.1128/msphere.00386-19.

8. Sidik, S.M., Huet, D., Ganesan, S.M., Huynh, M.H., Wang, T., Nasamu, A.S., Thiru, P., Saeij, J.P.J., Carruthers, V.B., Niles, J.C., et al. (2016). A Genome-wide CRISPR Screen in Toxoplasma Identifies Essential Apicomplexan Genes. Cell 166, 1423-1435.e12. 10.1016/j.cell.2016.08.019.

9. Li, W., Grech, J., Stortz, J.F., Gow, M., Periz, J., Meissner, M., and Jimenez-Ruiz, E. (2022). A splitCas9 phenotypic screen in Toxoplasma gondii identifies proteins involved in host cell egress and invasion. Nat. Microbiol. 7, 882–895. 10.1038/s41564-022-01114-y.

10. Benns, H.J., Storch, M., Falco, J.A., Fisher, F.R., Tamaki, F., Alves, E., Wincott, C.J., Milne, R., Wiedemar, N., Craven, G., et al. (2022). CRISPR-based oligo recombineering prioritizes apicomplexan cysteines for drug discovery. Nat. Microbiol. 7, 1891–1905. 10.1038/s41564-022-01249-y.

11. Shen, B., Brown, K.M., Lee, T.D., and Sibley, L.D. (2014). Efficient gene disruption in diverse strains of Toxoplasma gondii using CRISPR/CAS9. mBio 5, e01114–14. 10.1128/mBio.01114-14.

12. Hildebrandt, F. N., Matias, A., and Treeck, M. (2025). A CRISPR view on genetic screens in Toxoplasma gondii. Curr. Opin. Microbiol. 83, 102577. 10.1016/j.mib.2024.102577.

13. Young, J., Dominicus, C., Wagener, J., Butterworth, S., Ye, X., Kelly, G., Ordan, M., Saunders, B., Instrell, R., Howell, M., et al. (2019). A CRISPR platform for targeted in vivo screens identifies Toxoplasma gondii virulence factors in mice. Nat. Commun. 10, 3963. 10.1038/s41467-019-11855-w.

14. Ranganathan, V., Wahlin, K., Maruotti, J., and Zack, D.J. (2014). Expansion of the CRISPR–Cas9 genome targeting space through the use of H1 promoter-expressed guide RNAs. Nat. Commun. 5, 4516. 10.1038/ncomms5516.

15. Adamson, B., Norman, T.M., Jost, M., Cho, M.Y., Nuñez, J.K., Chen, Y., Villalta, J.E., Gilbert, L.A., Horlbeck, M.A., Hein, M.Y., et al. (2016). A Multiplexed Single-Cell CRISPR Screening Platform Enables Systematic Dissection of the Unfolded Protein Response. Cell 167, 1867-1882.e21. 10.1016/j.cell.2016.11.048.

16. Donald, R.G., and Roos, D.S. (1995). Insertional mutagenesis and marker rescue in a protozoan parasite: cloning of the uracil phosphoribosyltransferase locus from Toxoplasma gondii. Proc. Natl. Acad. Sci. U. S. A. 92, 5749–5753. 10.1073/pnas.92.12.5749.

17. Mefferd, A.L., Kornepati, A.V.R., Bogerd, H.P., Kennedy, E.M., and Cullen, B.R. (2015). Expression of CRISPR/Cas single guide RNAs using small tRNA promoters. RNA 21, 1683–1689. 10.1261/rna.051631.115.

18. Wei, Y., Qiu, Y., Chen, Y., Liu, G., Zhang, Y., Xu, L., and Ding, Q. (2016). CRISPR/Cas9 with single guide RNA expression driven by small tRNA promoters showed reduced editing efficiency compared to a U6 promoter. 10.1261/rna.057596.

19. Baer, M., Nilsen, T.W., Costigan, C., and Altman, S. (1990). Structure and transcription of a human gene for H1 RNA, the RNA component of human RNase P. Nucleic Acids Res. 18, 97–103. 10.1093/nar/18.1.97.

20. Hu, X., Meng, X., Li, J., Wang, K., and Yu, H. (2020). Improving the efficiency of the CRISPR-Cas12a system with tRNA-crRNA arrays. Crop J. 8, 403–407. 10.1016/j.cj.2019.06.007.

21. Mack, D., Mcleod, R., and Stark, B.C. (1996). Characterization of Ribonuclease P from Toxoplasma gondii.

22. Chon, C., Chon, G., Matsui, Y., Zeng, H., Lai, Z.-C., and Liu, A. (2021). Efficient multiplexed genome engineering with a polycistronic tRNA and CRISPR guide-RNA reveals an important role of detonator in reproduction of Drosophila melanogaster. PLoS One 16, e0245454..

23. Kozgunova, E. (2026). tRNA-based polycistronic CRISPR/Cas9 system boosts efficiency of multi-gene deletion in the moss Physcomitrella. Plant Methods 22, 28. 10.1186/s13007-026-01509-6.

24. Schindelin, J., Arganda-Carreras, I., Frise, E., Kaynig, V., Longair, M., Pietzsch, T., Preibisch, S., Rueden, C., Saalfeld, S., Schmid, B., et al. (2012). Fiji: An open-source platform for biological-image analysis. Nat. Methods 9, 676–682. 10.1038/nmeth.2019.

