## Supplementary Table 1 for "Using Polycistronic gRNAs and a tRNA Cleavage System in CRISPR-Based Genome Editing of *Toxoplasma gondii*"

| Construct | Construct Sequences |
| --- | --- |
| H1-SAG1gRNA-cr2 | GCGAGTTGAAATCGTCGTGGGGACGATTGCACCGCGGCCACATATTTAGCATGTCGCTATGTGTTCTGGGAAACTTGACCTAAGTGTAAAGTTGAGATTTCCTTCAGGTTTATATAGTTCTGTATGAGATCCACTCTTTCCCGGCAGTGAGACGCGCCGTCAGTTTGAGAGCTAAGCAGAAAGCTGCATAGCAAGTTCAAATAAGGCTAGTCCGTACACAACTTGAAAAAGTGGCAGCCGAGTCGGCTGCTTTTTTTTGAGCTCCAGCTTTTGTTCCCTTTAGTGAGGGTTAATTTCGAG |
| H1-SAG1gRNA-NORMtracr | CAAAGTGCGCGAGTTGAAATCGTCGTGGGGACGATTGCACCGCGGCCACATATTTAGCATGTCGCTATGTGTTCTGGGAAACTTGACCTAAGTGTAAAGTTGAGATTTCCTTCAGGTTTATATAGTTCTGTATGAGATCCACTCTTTCCCGGCAGTGAGACGCGCCGTCAGTTTTAGAGCTAGAAATAGCAAGTTAAAATAAGGCTAGTCCGTTATCAACTTGAAAAGTGGCACCGAGTCGGTGCTTTTTTGAGCTCCAGCTTTTGTTCCCTTTAGTGAGGGTTAATTTCGAGCTTGGCG |
| H1-SAG1gRNA-NORMtracr-TtRNA-UPRTgRNA-cr2 | TCTGGTGGGTCTCTGTCCATATTTAGCATGTCGCTATGTGTTCTGGGAAACTTGACCTAAGTGTAAAGTTGAGATTTCCTTCAGGTTTATATAGTTCTGTATGAGATCCACTCTTTCCCGGCAGTGAGACGCGCCGTCAGTTTTAGAGCTAGAAATAGCAAGTTAAAATAAGGCTAGTCCGTTATCAACTTGAAAAAGTGGCACCGAGTCGGTGCTTTTTTACGCTGTCAAGCGTTTGTGGTCTAGTGGTAGAATACCTCGTTGCCATCGAGGTGACCCGGGTTCGATTCCCGGCAGACGCAGGGCGTCTCGATTGTGAGAGCGTTTGAGAGCTAAGCAGAAAGCTGCATAGCAAGTTCAAATAAGGCTAGTCCGTACACAACTTGAAAAAGTGGCAGCCGAGTCGGCTGCTTTTTTTTGGCTCGGGAGACCTATCG |
| H1-UPRTgRNA-NORMtracr-TtRNA-SAG1gRNA-cr2 | TCTGGTGGGTCTCTGTCCATATTTAGCATGTCGCTATGTGTTCTGGGAAACTTGACCTAAGTGTAAAGTTGAGATTTCCTTCAGGTTTATATAGTTCTGTATGAGATCCACTCTTTCCCGGCGTCTCGATTGTGAGAGCGTTTTAGAGCTAGAAATAGCAAGTTAAAATAAGGCTAGTCCGTTATCAACTTGAAAAAGTGGCACCGAGTCGGTGCTTTTTTACGCTGTCAAGCGTTTGTGGTCTAGTGGTAGAATACCTCGTTGCCATCGAGGTGACCCGGGTTCGATTCCCGGCAGACGCAGGGCAGTGAGACGCGCCGTCAGTTTGAGAGCTAAGCAGAAAGCTGCATAGCAAGTTCAAATAAGGCTAGTCCGTACACAACTTGAAAAAGTGGCAGCCGAGTCGGCTGCTTTTTTTTGGCTCGGGAGACCTATCG |
| U6-SAG1gRNA-cr2 | ATGTTGGAGACACTGAGGACACACGGGAAACGCGAAAGATTTCAAATTAACGTACCCAAGCGCGAAAGCTTGCGCAGCATACACTCGAAGCGAACATCCCGAACCATCGAGAGGCGGAGAGCGATAAGTCTTTCACGCTGCGAAGTGTTGCGACGGCTGCGCCGCTGCACTGTGAATTGGGCGCCAATATTGCATCCTAGGCCTGACGCGCCTCCTGCAGAACGCGAGACACTGGGATATGTAGAGCCAAGGGGGAAACCTTCGAACTCTCGAATGTCTTCTCTGACAAGAATCATATTTCCATCAGTTCTGTCAGATTTTCAAATGGCGACCTGCAGAGGCCTGCTTCCTCCCTGTGCGCTCTTCGAAGGGGCTTTCTGTCGCGCAGGGTCACCTCGTCCCCGAAGGGGGTGTTTGCCTTCTGGTAAATGGGGATGTCAAGTTGGCAGTGAGACGCGCCGTCAGTTTGAGAGCTAAGCAGAAAGCTGCATAGCAAGTTCAAATAAGGCTAGTCCGTACACAACTTGAAAAAGTGGCAGCCGAGTCGGCTGCTTTTTTTTGAGCTCCAGCTTTTGTTC |
| U6-SAG1gRNA-NORMtracr | ATGTTGGAGACACTGAGGACACACGGGAAACGCGAAAGATTTCAAATTAACGTACCCAAGCGCGAAAGCTTGCGCAGCATACACTCGAAGCGAACATCCCGAACCATCGAGAGGCGGAGAGCGATAAGTCTTTCACGCTGCGAAGTGTTGCGACGGCTGCGCCGCTGCACTGTGAATTGGGCGCCAATATTGCATCCTAGGCCTGACGCGCCTCCTGCAGAACGCGAGACACTGGGATATGTAGAGCCAAGGGGGAAACCTTCGAACTCTCGAATGTCTTCTCTGACAAGAATCATATTTCCATCAGTTCTGTCAGATTTTCAAATGGCGACCTGCAGAGGCCTGCTTCCTCCCTGTGCGCTCTTCGAAGGGGCTTTCTGTCGCGCAGGGTCACCTCGTCCCCGAAGGGGGTGTTTGCCTTCTGGTAAATGGGGATGTCAAGTTGGCAGTGAGACGCGCCGTCAGTTTTAGAGCTAGAAATAGCAAGTTAAAATAAGGCTAGTCCGTTATCAACTTGAAAAGTGGCACCGAGTCGGTGCTTTTTTGAGCTCCAGCTTTTGTTC |
| U6-SAG1gRNA-NORMtracr-HtRNA-UPRTgRNA-cr2 | TCTGGTGGGTCTCTGTCCATGTTGGAGACACTGAGGACACACGGGAAACGCGAAAGATTTCAAATTAACGTACCCAAGCGCGAAAGCTTGCGCAGCATACACTCGAAGCGAACATCCCGAACCATCGAGAGGCGGAGAGCGATAAGTCTTTCACGCTGCGAAGTGTTGCGACGGCTGCGCCGCTGCACTGTGAATTGGGCGCCAATATTGCATCCTAGGCCTGACGCGCCTCCTGCAGAACGCGAGACACTGGGATATGTAGAGCCAAGGGGGAAACCTTCGAACTCTCGAATGTCTTCTCTGACAAGAATCATATTTCCATCAGTTCTGTCAGATTTTCAAATGGCGACCTGCAGAGGCCTGCTTCCTCCCTGTGCGCTCTTCGAAGGGGCTTTCTGTCGCGCAGGGTCACCTCGTCCCCGAAGGGGGTGTTTGCCTTCTGGTAAATGGGGATGTCAAGTTGGCAGTGAGACGCGCCGTCAGTTTTAGAGCTAGAAATAGCAAGTTAAAATAAGGCTAGTCCGTTATCAACTTGAAAAAGTGGCACCGAGTCGGTGCTTTTTTAACAAAGCACCAGTGGTCTAGTGGTAGAATAGTACCCTGCCACGGTACAGACCCGGGTTCGATTCCCGGCTGGTGCAGGCGTCTCGATTGTGAGAGCGTTTGAGAGCTAAGCAGAAAGCTGCATAGCAAGTTCAAATAAGGCTAGTCCGTACACAACTTGAAAAAGTGGCAGCCGAGTCGGCTGCTTTTTTTTGGCTCGGGAGACCTATCG |
| U6-SAG1gRNA-NORMtracr-TtRNA-UPRTgRNA-cr2 | TCTGGTGGGTCTCTGTCCATGTTGGAGACACTGAGGACACACGGGAAACGCGAAAGATTTCAAATTAACGTACCCAAGCGCGAAAGCTTGCGCAGCATACACTCGAAGCGAACATCCCGAACCATCGAGAGGCGGAGAGCGATAAGTCTTTCACGCTGCGAAGTGTTGCGACGGCTGCGCCGCTGCACTGTGAATTGGGCGCCAATATTGCATCCTAGGCCTGACGCGCCTCCTGCAGAACGCGAGACACTGGGATATGTAGAGCCAAGGGGGAAACCTTCGAACTCTCGAATGTCTTCTCTGACAAGAATCATATTTCCATCAGTTCTGTCAGATTTTCAAATGGCGACCTGCAGAGGCCTGCTTCCTCCCTGTGCGCTCTTCGAAGGGGCTTTCTGTCGCGCAGGGTCACCTCGTCCCCGAAGGGGGTGTTTGCCTTCTGGTAAATGGGGATGTCAAGTTGGCAGTGAGACGCGCCGTCAGTTTTAGAGCTAGAAATAGCAAGTTAAAATAAGGCTAGTCCGTTATCAACTTGAAAAAGTGGCACCGAGTCGGTGCTTTTTTACGCTGTCAAGCGTTTGTGGTCTAGTGGTAGAATACCTCGTTGCCATCGAGGTGACCCGGGTTCGATTCCCGGCAGACGCAGGGCGTCTCGATTGTGAGAGCGTTTGAGAGCTAAGCAGAAAGCTGCATAGCAAGTTCAAATAAGGCTAGTCCGTACACAACTTGAAAAAGTGGCAGCCGAGTCGGCTGCTTTTTTTTGGCTCGGGAGACCTATCG |
| U6-UPRTgRNA-NORMtracr-HtRNA-SAG1gRNA-cr2 | TCTGGTGGGTCTCTGTCCATGTTGGAGACACTGAGGACACACGGGAAACGCGAAAGATTTCAAATTAACGTACCCAAGCGCGAAAGCTTGCGCAGCATACACTCGAAGCGAACATCCCGAACCATCGAGAGGCGGAGAGCGATAAGTCTTTCACGCTGCGAAGTGTTGCGACGGCTGCGCCGCTGCACTGTGAATTGGGCGCCAATATTGCATCCTAGGCCTGACGCGCCTCCTGCAGAACGCGAGACACTGGGATATGTAGAGCCAAGGGGGAAACCTTCGAACTCTCGAATGTCTTCTCTGACAAGAATCATATTTCCATCAGTTCTGTCAGATTTTCAAATGGCGACCTGCAGAGGCCTGCTTCCTCCCTGTGCGCTCTTCGAAGGGGCTTTCTGTCGCGCAGGGTCACCTCGTCCCCGAAGGGGGTGTTTGCCTTCTGGTAAATGGGGATGTCAAGTTGGCGTCTCGATTGTGAGAGCGTTTTAGAGCTAGAAATAGCAAGTTAAAATAAGGCTAGTCCGTTATCAACTTGAAAAAGTGGCACCGAGTCGGTGCTTTTTTAACAAAGCACCAGTGGTCTAGTGGTAGAATAGTACCCTGCCACGGTACAGACCCGGGTTCGATTCCCGGCTGGTGCAGGCAGTGAGACGCGCCGTCAGTTTGAGAGCTAAGCAGAAAGCTGCATAGCAAGTTCAAATAAGGCTAGTCCGTACACAACTTGAAAAAGTGGCAGCCGAGTCGGCTGCTTTTTTTTGGCTCGGGAGACCTATCG |
| U6-UPRTgRNA-NORMtracr-TtRNA-SAG1gRNA-cr2 | TCTGGTGGGTCTCTGTCCATGTTGGAGACACTGAGGACACACGGGAAACGCGAAAGATTTCAAATTAACGTACCCAAGCGCGAAAGCTTGCGCAGCATACACTCGAAGCGAACATCCCGAACCATCGAGAGGCGGAGAGCGATAAGTCTTTCACGCTGCGAAGTGTTGCGACGGCTGCGCCGCTGCACTGTGAATTGGGCGCCAATATTGCATCCTAGGCCTGACGCGCCTCCTGCAGAACGCGAGACACTGGGATATGTAGAGCCAAGGGGGAAACCTTCGAACTCTCGAATGTCTTCTCTGACAAGAATCATATTTCCATCAGTTCTGTCAGATTTTCAAATGGCGACCTGCAGAGGCCTGCTTCCTCCCTGTGCGCTCTTCGAAGGGGCTTTCTGTCGCGCAGGGTCACCTCGTCCCCGAAGGGGGTGTTTGCCTTCTGGTAAATGGGGATGTCAAGTTGGCGTCTCGATTGTGAGAGCGTTTTAGAGCTAGAAATAGCAAGTTAAAATAAGGCTAGTCCGTTATCAACTTGAAAAAGTGGCACCGAGTCGGTGCTTTTTTACGCTGTCAAGCGTTTGTGGTCTAGTGGTAGAATACCTCGTTGCCATCGAGGTGACCCGGGTTCGATTCCCGGCAGACGCAGGGCAGTGAGACGCGCCGTCAGTTTGAGAGCTAAGCAGAAAGCTGCATAGCAAGTTCAAATAAGGCTAGTCCGTACACAACTTGAAAAAGTGGCAGCCGAGTCGGCTGCTTTTTTTTGGCTCGGGAGACCTATCG |
